# *RPE65* knockout *Xenopus laevis* have a compromised but detectable electroretinogram and altered visual responses, without retinal degeneration or altered melanophore dispersion

**DOI:** 10.1101/2025.08.15.670600

**Authors:** Colette N. Chiu, Beatrice M. Tam, Paloma Burns, Orson L. Moritz

## Abstract

The isomerohydrolase RPE65 is a critical element of the visual cycle, the series of enzymatic reactions by which the chromophore of the visual pigments is regenerated following light exposure. In humans, mutations in the *rpe65* gene cause a severe form of blindness called Leber’s congenital amaurosis. Studies of *RPE65^-/-^* mice have shown dramatic depletion of 11- cis-retinal in the retina, resulting in a slow retinal degeneration. However, a number of studies suggest that RPE65 may not be necessary for the regeneration of photopigment in all photoreceptor types. Using CRISPR/Cas9 technology, we previously generated *RPE65* knockout *Xenopus laevis* in order to test the involvement of rhodopsin chromophore in the cell death mechanisms associated with *rhodopsin* mutations and rhodopsin quality control. Here we further characterize the effects of *RPE65* knockout in these animals, and show their rod photoreceptors have shortened outer segments that lack detectable rhodopsin photopigment. However, there is no progressive degeneration of rods or cones. Via electroretinography we found greatly reduced but significant responses to light under scotopic and photopic conditions. We also found reduced behavioral sensitivity to light, while light-induced melanophore dispersion was unaffected. *RPE65* knockout *X. laevis* may be a useful system for examining RPE65-independent photosensation mechanisms in vertebrates.

## 1. Introduction

G-protein coupled receptor-based visual pigments such as rhodopsins and photopsins utilize 11- cis retinal or closely related vitamin-A derivatives as a covalently-bound chromophore. On absorption of a photon, the bound chromophore is isomerized from the 11-cis form to the all-trans form, promoting an altered protein conformation and initiating the phototransduction cascade (Molday and Moritz, 2015). Subsequently, all-trans retinal spontaneously dissociates from the receptor protein, and a new molecule of 11-cis retinal binds, regenerating the visual pigment. 11-cis retinal is regenerated from all-trans retinal via the visual cycle, a complex set of reactions largely carried out in the closely associated retinal pigment epithelium (Saari, 2000). The critical conversion from all-trans to 11-cis is carried out by the enzyme RPE65, which converts all-*trans*-retinyl ester to 11-*cis*-retinol, which is then oxidized to 11-*cis*-retinal, the ligand for rod and cone opsins, by downstream enzymes (Cai et al., 2009; Jin et al., 2005; Moiseyev et al., 2005; Redmond et al., 2005, 1998).

Loss of function of the gene encoding RPE65 causes a severe form of blindness called Leber’s congenital amaurosis (LCA) in humans (Marlhens et al., 1997), and similar visual disfunction in mice and dogs (Redmond et al., 1998; Veske et al., 1999). Patients with RPE65-associated LCA have very little visual capability. In contrast to disorders such as retinitis pigmentosa or macular degeneration, this is largely due to loss of function rather than neurodegeneration (although degeneration also occurs), making these individuals good candidates for gene therapy (Acland et al., 2005, 2001; Bainbridge et al., 2015; Cideciyan et al., 2008). Interestingly, LCA patients do retain some light sensitivity, likely mediated by cones, suggesting that their retinas contain other pathways for photopigment regeneration (Jacobson et al., 2009).

Other studies have proposed the presence of alternative pathways for pigment generation (Wang and Kefalov, 2011). Among the earliest evidence was the observation that frog long-wavelength sensitive (LWS) cone and green rod photopigments can regenerate in the absence of the RPE (Bruch Goldstein and Wolf, 1973; Goldstein, 1970). More recent studies have proposed pathways present in Muller glia, potentially involving an alternative isomerase DES1 (Kaylor et al., 2013), and the retinyl-ester synthase MFAT (Kaylor et al., 2014), although the involvement of Muller cell-derived DES1 has more recently been disproven (Kiser et al., 2019; Ward et al., 2020). Alternatively, these pathways may utilize photoconversion by blue light, with the involvement of catalysts such as phospholipids or RGR opsin (Chen et al., 2001; Kaylor et al., 2017; Morshedian et al., 2019; Tworak et al., 2023). Similar pathways for chromophore regeneration by photoisomerization exist in the rhabdomeric photoreceptors of non-vertebrates such as insects, although these species utilize bistable photopigments from which the chromophore does not dissociate (Montell, 2012; Stavenga, 1995). There is also evidence that 9- cis retinal can act as an alternative chromophore in the absence of RPE65, although its source is unclear (Fan et al., 2003).

Here we examine the effect of *RPE65* knockout on retinal morphology and function in a non-mammalian vertebrate, *Xenopus laevis* (African clawed frog). Notably, the first evidence for alternative chromophore regeneration pathways was obtained from frog retina (Bruch Goldstein and Wolf, 1973; Goldstein, 1970) and the *Xenopus laevis* genome encodes 22 different type II opsins (not including gene duplicates) including visual opsins, non-visual opsins, pineal opsins, neuropsins, photoisomerases and melanopsins (Bertolesi et al., 2022), making it a versatile system for examining these processes. We generated RPE65 knockouts using CRISPR/Cas9 technology, and determined the effects on the retina via histology, *in-vivo* imaging, difference spectroscopy, and electroretinography, as well as assays of visual and non-visual responses to light. We also examined light-driven responses of melanophores. Our results indicate the retinas of these animals cannot generate detectable amounts of rhodopsin in the absence of RPE65. However, we found relatively robust responses of the retina and melanosomes to light that do not require RPE65.

## 2. Methods

### 2.1 Breeding

Generation of F0 RPE65 animals was previously described (Tam et al., 2024). The animals from this study were raised to maturity. These animals were crossed to WT, and the resulting embryos were analyzed by Sanger sequencing as previously described (Feehan et al., 2017; Ghaseminejad et al., 2022). A male and female that transmitted high proportions of knockout alleles in *RPE65.L* and *RPE65.S* were chosen and mated against each other. The resulting F1 animals were screened by Sanger sequencing and raised to maturity. A pair of animals carrying only out of frame edits in both *RPE65.L* and *RPE65.S* (Table 1) were mated against each other to generate full knock out animals in the F2 generation.

### 2.2 Western Blot

Normally developed tadpoles were sacrificed at 14dpf (developmental stage 48). One eye of each tadpole was solubilized in 100µl of a 1:1 mixture of phosphate-buffered saline and SDS-PAGE loading buffer containing 1mM EDTA and 100µg/ml PMSF. Samples were separated on 10% SDS-PAGE gels and transferred to Immobilon-P membranes. Membranes were probed with 1:3000 rabbit polyclonal anti- RPE65 (YZ5437, Wen et al., 2019) overnight, followed by 1:5000 IR-dye800-conjugated goat anti-rabbit secondary antibody (LI-COR Biosciences). A LI-COR Odyssey imaging system was used to image and quantify the integrated intensity of blot signals. The membrane was subsequently re-probed with 1:20 anti-rod opsin mAb B630N, followed by 1:10000 IR-dye800-conjugated goat anti-mouse secondary antibody (LI-COR Biosciences).

### 2.3 Optical Coherence Tomography

Adult frogs, 2 years old, WT or *RPE65* KO, were anesthetized using 0.1% buffered tricaine solution. A drop of clear soft eye gel was placed on the frog eye before imaging using Phoenix Research Systems Micron V fundus camera and OCT scan head.

### 2.4 Confocal Microscopy

Samples were prepared for confocal microscopy as previously described (Tam et al., 2024). One eye of each tadpole was fixed in 4% buffered paraformaldehyde, pH 7.4. Fixed eyes were infiltrated with 20% buffered sucrose for 2-6h, embedded in Optical Cutting Temperature Compound (OCT; Fisher HealthCare), and frozen. 12µm sections were cut using a cryostat, and labelled with 1:3000 anti-RPE65 (YZ5437, Wen et al., 2019), 1:500 anti-LWS (YZ3526), 1:1000 anti-SW1 (YZ7140), or1:300 anti-SWS2 (gift of Dr. M. Kono), followed by Cy3-conjugated anti-mouse or anti-rabbit secondary antibody (Jackson ImmunoResearch Laboratories) and Alexa Fluor 488-conjugated wheat germ agglutinin (WGA) (ThermoFisher Scientific) and Hoechst 33342 (Sigma Aldrich) counterstains. YZ7140 was raised against the *X. laevis SWS1*-derived peptide sequence GRPMSDDSSVSSTSQRT.

The labelled sections were imaged using Zeiss 880 Airyscan microscope with a 40x NA 1.2 water-immersion objective (Carl Zeiss). Zen software was used to generate maximum intesnisty projections, and to measure rod photoreceptor length in Figure 1D. Final figures were assembled using Adobe Photoshop software.

**Figure 1:**
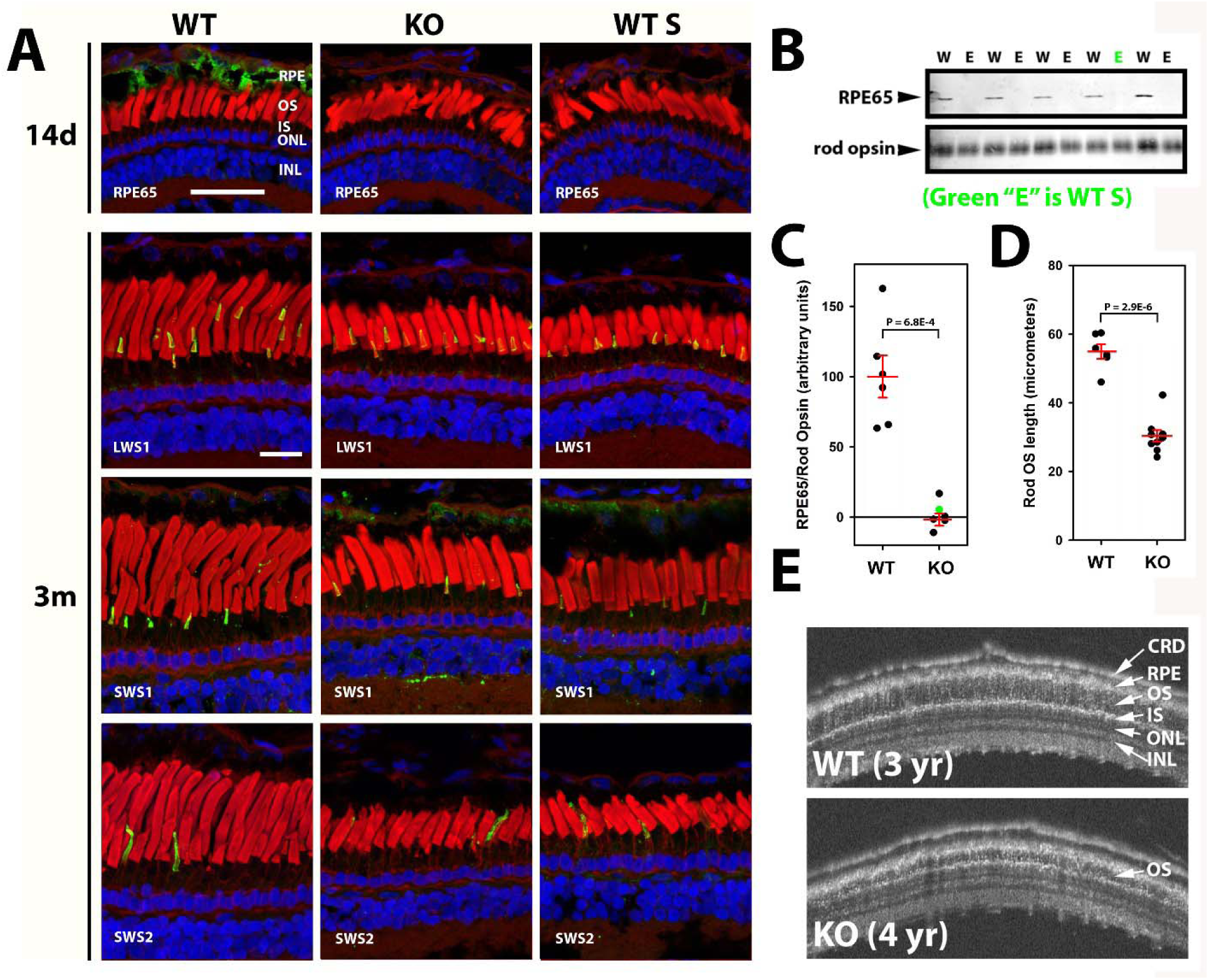
*RPE65* knockout X. laevis have no retinal degeneration, no RPE65 expression, and altered ROS length. **A:** Confocal microscopy of retinas from 14 d and 3m old animals labeled with antibodies shown (green) and counterstained with WGA (red) and Hoechst 33342 (blue), animals are wildtype, knockout, or have one WT “S” allele. **B:** western blot showing lack of RPE65 expression in edited animals, including one sample from an animal with one WT “S” allele. W – WT, E = edited (WT S is “E” labeled in green). **C:** quantification of western blot data shown in B. **D:** Quantification of OS length shown in A. In C and D, the average value and +/- S.E.M. are shown in red. **E:** OCT imaging of an older knockout animal (4 years) compared to WT. The OS layer is still intact.

### 2.5 Difference Spectroscopy

Post-metamorphic froglets (3-4 months old) were dark adapted overnight. Both eyes were dissected in the dark under dim red light and solubilized in 100µl detergent containing 1mM EDTA, 1mM PMSF, 1% octyl-β- D-glucopyranoside, and 10mM Tris pH 7.4. The solubilized eyes were centrifuged for 1 hour at 55000rpm at 4C using TLA55 rotor in an Optima Max E Ultracentrifuge. The absorbance spectrum of the supernatant (200-700nm) was obtained using a DU730 UV/Vis Spectrophotometer (Beckman Coulter). The sample was then bleached with bright white light for 30 seconds, and another spectrum was taken; the difference spectra was obtained by subtraction.

### 2.6 Electroretinography

Electroretinography was performed as previously described (Vent-Schmidt et al., 2017). Animals aged 6 weeks (developmental stage 53) or 3 months (post metamorphic froglets) were anesthetized using 0.01% or 0.1% buffered tricaine. The animals were placed on a modified petri dish containing an ECG electrode, and a glass recording electrode was placed onto the cornea with a drop of artificial tears. Scotopic or photopic electroretinograms of an increasing series of blue or white flashes, respectively, were collected using the Espion Ganzfeld electroretinography unit (Diagnosys) (see results for details of individual protocols). Spectral sensitivities under scotopic and photopic conditions were determined using flashes generated from LEDs of 448nm, 530nm, 590nm, and 627nm, while holding the flash power constant. 4-7 animals were analyzed for each condition.

### 2.7 Visual Behavior assay

Groups of 20 tadpoles stage 48 (14dpf), either WT or *RPE65* KO, were placed in a 4-liter plastic tank divided into two equal sections, one colored white and one colored black. The tank was placed on a bench lit from above with standard white fluorescent room lighting, measured at 450 lux. The tadpoles were acclimatized to the testing bin for 5min. The tank was photographed four times at one-minute intervals, rotated 180 degrees, and again photographed four times at one-minute intervals. The number of tadpoles on the white and black backgrounds were determined for each time point, and averaged across all eight observations. For each genotype, 5 groups of 20 animals were similarly assayed 8 times. The average assay result for each group was calculated, resulting in n = 5 data points for each genotype, with each data point based on 160 total observations.

### 2.8 Pigment Dispersion Assay

*X. laevis* larvae (stage 44, 6dpf) were anesthetized in 0.01% buffered tricaine for 0.5-1 minute before eye enucleation with a microblade. The larvae were placed back in ringer solution for recovery for 24 hours. Tadpoles at developmental stage 45 (7dpf) were dark adapted overnight. Under dim red light, tadpoles were anesthetized in 0.01% buffered tricaine for 1 min and transferred to a petri dish filled with agarose containing 0.01% buffered tricaine. The tadpoles were exposed to 800 lux white light on a Leica MZ16F microscope stage, and imaged using the microscope camera. Image files were analyzed using ImageJ to measure the total area of melanophore migration.

## 3. Results

### 3.1 Generation of genetically modified *X. laevis*

We previously determined that *X. laevis* have two genes encoding RPE65, *RPE65.S* and *RPE65.L*, and described methods for ablating these genes in F0 animals using CRISPR/Cas9- based gene editing (Feehan et al., 2019, 2017; Tam et al., 2024; Wen et al., 2019). We have now reared chimeric founder animals derived from these previous studies to maturity, and crossed them over two generations to generate animals that consistently transmit frameshifting mutations in all four *RPE65* alleles (Supplementary Table 1) (see **2.1** above for details of breeding). These alleles are predicted to generate mRNAs that are degraded by the nonsense-mediated decay pathway, resulting in loss of function.

### 3.2 Western blot and Histology

To determine whether our genetic manipulations eliminated expression of the RPE65 protein, we raised wildtype (WT) and edited animals to 14 days post fertilization, and processed eyes for histology and western blot analysis. We were unable to detect RPE65 protein by immunolabeling in animals carrying frame-shifting mutations in all four *RPE65* alleles (Figure 1A). Similarly, we were unable to detect RPE65 protein via western blot in the retinas of these animals (Figure 1B, C). We found no evidence of low-level expression in cones or other cells of the retina previously reported by some researchers (Ma et al., 1998; Znoiko et al., 2002) (Figure 1A). Some animals that we examined had one remaining wildtype allele in the *RPE65.S* gene. The RPE65 protein was also undetectable in the eyes of these animals by both immunolabeling and western blot (Figure 1A, B, C). Because the encoded epitope recognized by our antibody is identical between *RPE65.L* and *RPE65.S*, we concluded that the *RPE65.S* gene does not function in *X. laevis* retina at this stage of development. However, because the coding sequence of *RPE65.S* is consistent with a functional gene (i.e. there are no premature stop codons), it is possible that *RPE65.S* functions at a different stage of development, or in non-ocular cell types. This result is consistent with RNA seq developmental stage and tissue expression data available on the Xenbase website (Session et al., 2016) (https://www.xenbase.org/xenbase/), which indicate *RPE65.S* is expressed maximally at developmental stage 15 (neurula), with no expression in any tissue examined at later stages. For the experiments described below, we used animals with frameshifting mutations in all four *RPE65* alleles, which we refer to as “*RPE65* knockouts” or “*RPE65* KOs”, except where otherwise noted.

### 3.3 Retinal morphology

*RPE65* knockout animals examined at 3 months post-fertilization (stage 66, post-metamorphosis) had retinas with shorter rod photoreceptor outer segments relative to WT animals (Figure 1A, D). Outer segment length averaged 55 µm in WT and 30 µm in knockouts at age 3 months; in contrast, there was little or no difference at age 14 days post-fertilization (stage 49) (Figure 1A, D). In mice, *RPE65* knockout results in rapid degeneration of SWS cone photoreceptors, which is associated with mis-localization of the cone opsin protein to inner segments (Rohrer et al., 2005). Therefore, we examined LWS and SWS1 cones, as well as green rods (which express the *X. laevis* SWS2 opsin) in these animals by labeling with antibodies specific for each type of cone opsin. However, we found no evidence of cell loss or significant morphological changes (Figure 1A). Animals with one remaining WT *RPE65.S* allele similarly had shorter rod OS, and preservation of cones and green rods, a phenotype indistinguishable from knockouts (Figure 1A). Examination of older knockout animals (up to 4 years) by optical coherence tomography (Figure 1E) showed no additional abnormalities beyond shortened OS length.

### 3.4 Difference spectroscopy

To determine whether visual pigments were absent from the retinas of *RPE65* knockout *X. laevis*, we sacrificed animals held in darkness overnight, dissected eyes under dim red light, solubilized the eyes, and analyzed the soluble fractions by difference spectroscopy. Spectra were collected before and after exposure of the soluble fractions to bright white light, and difference spectra were calculated (Figure 2A). Using these conditions, we obtained a strong signal with a peak absorbance at 525 nm from WT eyes corresponding to rhodopsin, but no significant signal at 525 nm from *RPE65* knockout eyes, indicating lack of rhodopsin. Three smaller peaks at 415 nm, 540 nm, and 575 nm were also present. However, these do not match absorbance maxima of known *X. laevis* visual pigments (Bertolesi et al., 2022), and instead are consistent with the absorbance spectra of hemoglobin, i.e. contamination of the sample with blood.

**Figure 2:**
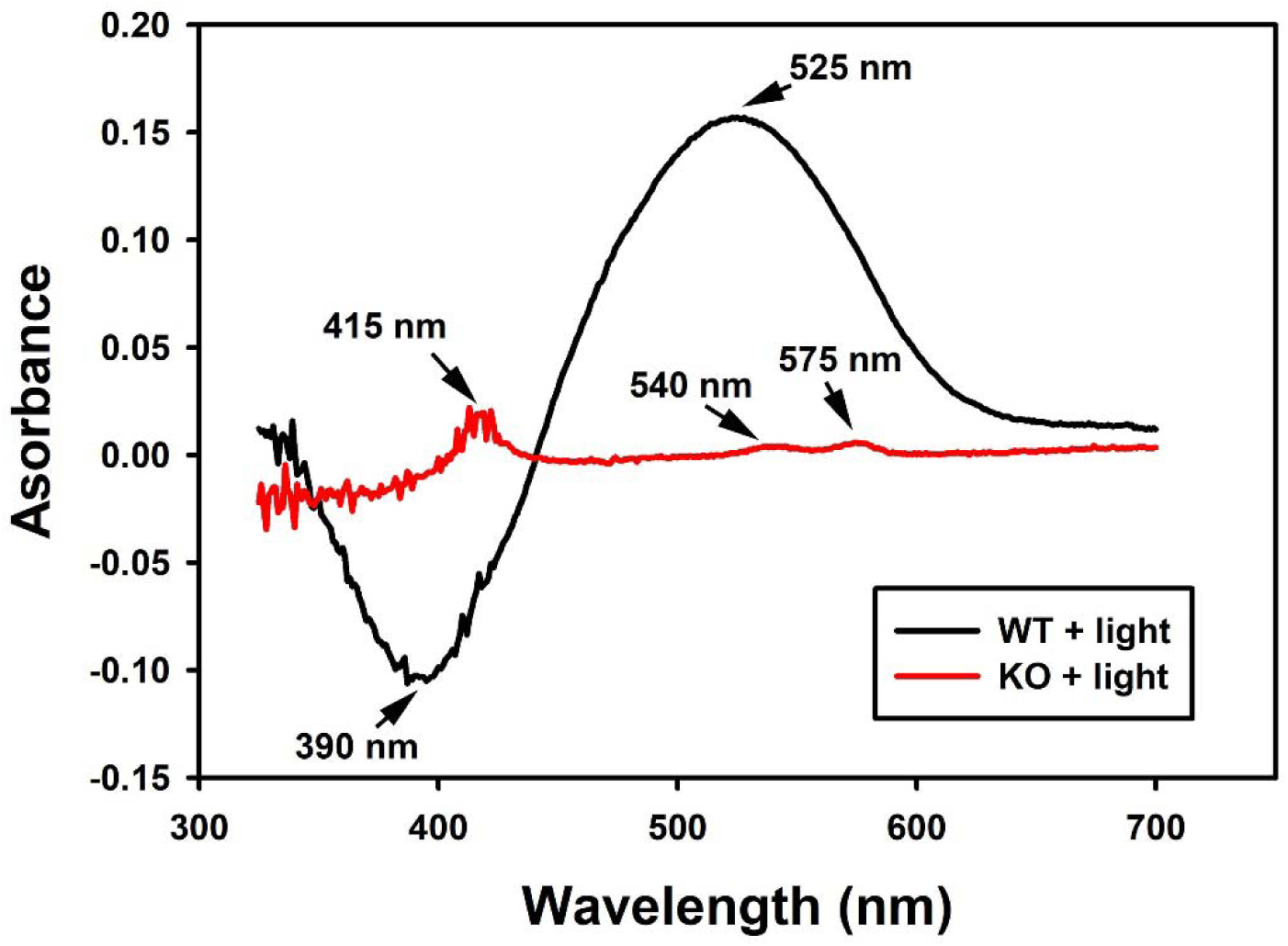
Difference spectra of WT vs KO animals. Black trace: sample obtained from a WT animal. Red trace: sample obtained from an RPE65 KO animal. Indicated peaks on the black trace are consistent with rhodopsin (525 nm) and meta-II rhodopsin (390 nm). Indicated peaks on the red trace are consistent with the spectra of hemoglobin.

### 3.5 Electroretinography

We investigated rod function in *RPE65* knockout animals by examining scotopic ERG responses, using a series of blue flashes of increasing intensity in dark-reared animals. As ERG signals become more robust as animals age, we examined animals at both 6 weeks post fertilization (stage 54-56) (Fig 3 A, B) and 3 months post fertilization (after metamorphosis) (Fig 3 C, D). In contrast to results previously obtained with *RPE65* KO mice, we noted significant ERG responses in our *RPE65* KO *X. laevis*. These responses were both reduced and delayed relative to WT responses, with a shift in sensitivity of approximately 2 orders of magnitude in both age groups. Overall, the responses of 3m old *RPE65* KO animals closely resembled the responses of WT animals to much dimmer flashes in both magnitude and time course (for example, in Fig 3 panel B compare the *RPE65* KO response to a 2500 cds/m2 flash to the WT response to a 25 cds/m2 flash), suggesting the responses are driven by the same photoreceptors (i.e. rods) despite the lack of detectable rhodopsin.

**Figure 3:**
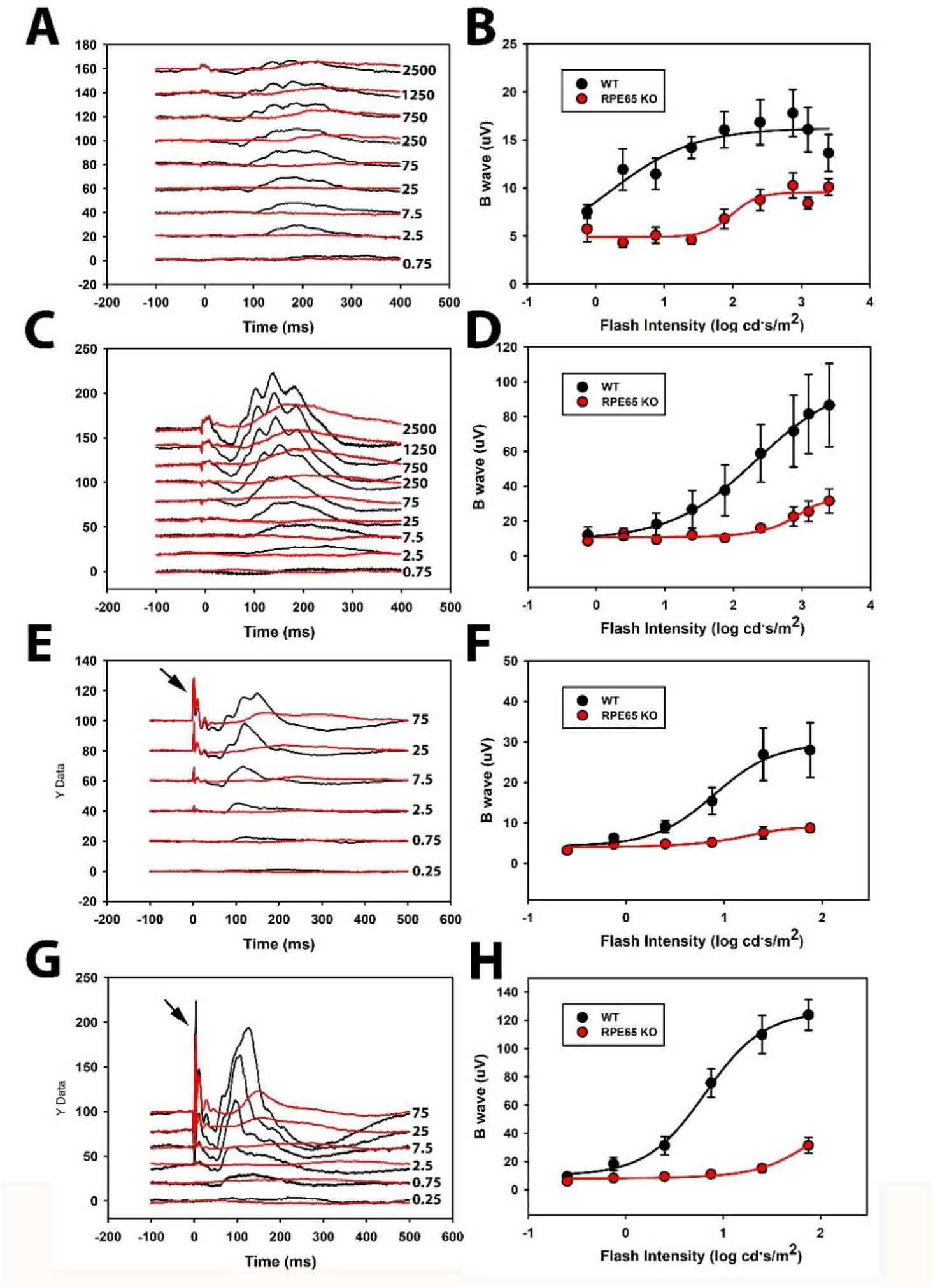
Electroretinography of WT and RPE65 KO *X. laevis*. Traces are shown in **A, C, E,** and **G**, and B wave amplitudes are graphed in **B, D, F** and **H. A-D** are scotopic ERGs on 6- week-old (**A, B**) or 3-month-old (**C, D**) animals. **E-H** are photopic ERGS for 6-week-old (**E-F**) or 3-month-old (**G-H**) animals. The black arrows indicate an artifact associated with the flash discharge. Flash intensities are indicated beside the traces, in cd*s/m^2^. Black traces or plots = WT, Red traces or plots = knockout. Error bars are +/- S.E.M. Traces represent averages of 5-7 animals; similarly, data points are averages of 5-7 animals.

We then investigated cone function in the *RPE65* KO animals by examining the photopic ERG responses to a series of white light flashes in the presence of a constant white background light in 6w and 3m old animals (Fig 3 panel E-G). We similarly observed detectable ERG responses, with a shift in sensitivity of roughly two orders of magnitude, and a significant delay in the responses. However, in this case, the delay of the ERG B wave in RPE65 KO animals was greater than that observed for any flash intensity in WT animals, suggesting the possibility of altered cone phototransduction and/or B-wave generation kinetics, or that an altered population of photoreceptors and bipolar cells generated the signal.

### 3.6 Spectral sensitivity

To further characterize these unusual ERG responses, we conducted several ERG protocols in which we varied the wavelength of the stimulus while holding the flash power constant, in order to generate an action spectrum. Flashes of 1.27E-05 watts were generated using LEDs emitting at 448nm, 530nm, 590nm, and 627nm; for the purpose of comparison with scotopic ERG conditions described above, this corresponded to an intensity of 12 cd.s/m2 for the blue light flash. To ensure that WT scotopic responses were not saturated, two sets of ERGs were obtained using two different flash intensities. For protocols conducted on dark adapted animals (i.e. scotopic conditions) (Fig 4 A-D), *RPE65* KO animals had reduced ERGs relative to WT animals at all four wavelengths. Another notable difference was that in WT animals the largest responses were to red flashes, suggesting that LWS cones contribute to these responses, while *RPE65* KO animals had largest responses to intermediate wavelengths. Similar results were obtained for light adapted animals examined in the presence of a constant background light (i.e. phototopic conditions) Fig 4 (E-H). Again, response size to all wavelengths was reduced. However, in this case the responses to red wavelengths were not differentially altered.

**Figure 4:**
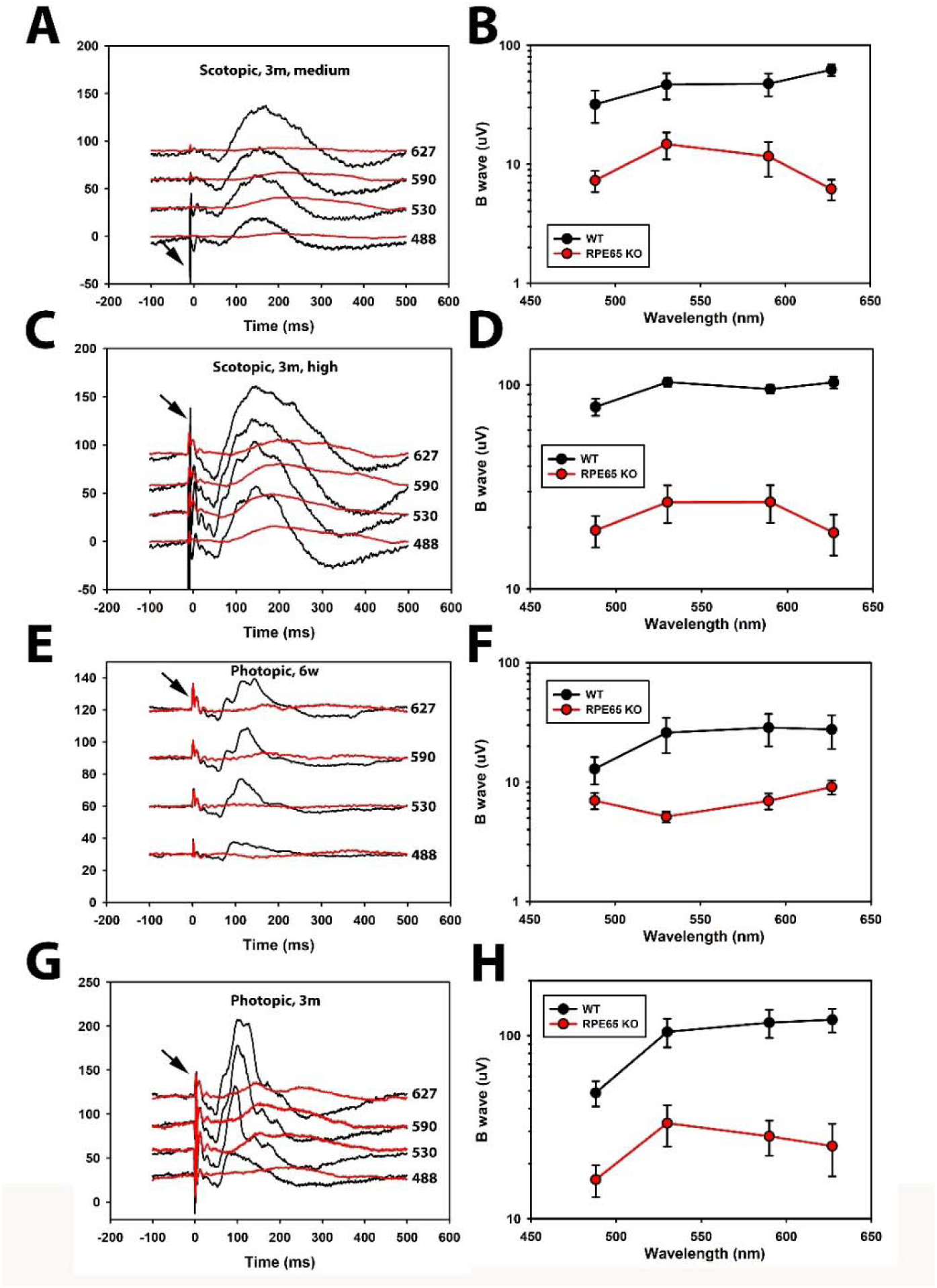
Spectral sensitivity of WT and RPE65 KO *X. laevis* investigated by electroretinography. Traces are shown in **A, C, E**, and **G**, and B wave amplitudes are graphed in **B, D, F** and **H. A-D** are scotopic ERGs at 4 different wavelengths, performed on 3-month-old wildtype (black) or knockout (red) animals at either medium (**A-B**) or high intensity (**C-D**) flashes. Flash wavelength is shown next to the traces. **E-H** are photopic ERGs performed on 6-week-old animals (**E-F**), or 3-month-old animals (**G-H**) performed at the same wavelengths as above. Error bars are +/- S.E.M. Traces represent averages of 4-7 animals; similarly, data points are averages of 5-7 animals.

### 3.7 Visual behaviors

At early developmental stages, *Xenopus laevis* tadpole eyes are directed downwards, and the animals show a strong preference for swimming above light-colored backgrounds (Moriya et al., 1996; Viczian and Zuber, 2014). To determine whether *RPE65* knockout affected this behavior, we assayed groups of 20 animals (either WT or *RPE65* KOs) as detailed under **2.7**. We found that WT animals chose the light background 75.6 +/- 0.9% of the time, while *RPE65* KO animals chose the light background 54.4 +/- 2.5% of the time (+/-95% confidence intervals) (Fig 5).

**Figure 5:**
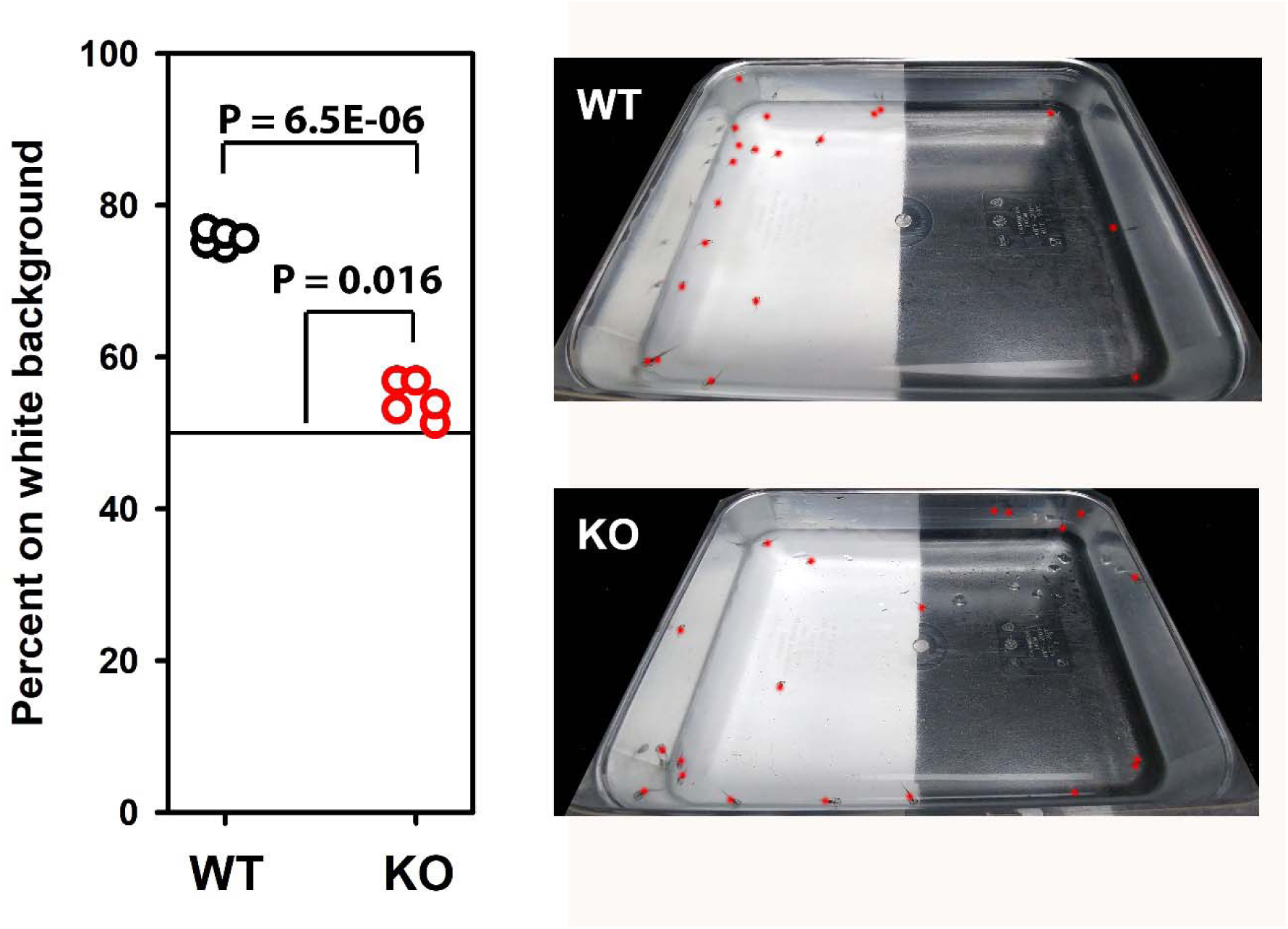
Visual function in tadpoles tested using a behavioral assay. Assay results are shown on the left, and representative images from the assay are shown on the right. In the images, the location of each tadpole is marked using a red dot. WT tadpoles prefer to swim over a light background. KO tadpoles show less preference, with a distribution close to random. P values are derived from 1 and 2-sample T tests.

Thus, this visual behavior was strongly inhibited in *RPE65* KO animals. Although the behavior of the knockouts was close to random, our results indicate *RPE65* KO animals likely retain a small preference for swimming over light backgrounds (P = 0.016 for one-sample T-test).

### 3.8 Melanophore responses to light

*X. laevis* tadpoles alter their pigmentation in response to both ambient lighting and the shade of the background over which they are swimming. This occurs by dispersion of melanophores in the melanocytes of the skin, causing a transition from a light to a darker appearance. The response to ambient light is driven by the pineal complex, while responses to the background originate in the eye (Bertolesi et al., 2020). We examined the dependence of melanophore dispersion in *X. laevis* tadpoles on light in wildtype and *RPE65* KO animals, by transferring dark adapted animals to light conditions (an illuminated dissecting microscope stage), and quantifying the pigmented area via micrographs taken at regular intervals, as detailed under **2.8**. In addition, we examined the dependence of these results on the presence of eyes, by examining animals that were enucleated at an early stage of development (stage 44) (Fig 6). Results were analyzed by 3- way ANOVA for dependence on time, enucleation, and *RPE65* genotype.

**Figure 6:**
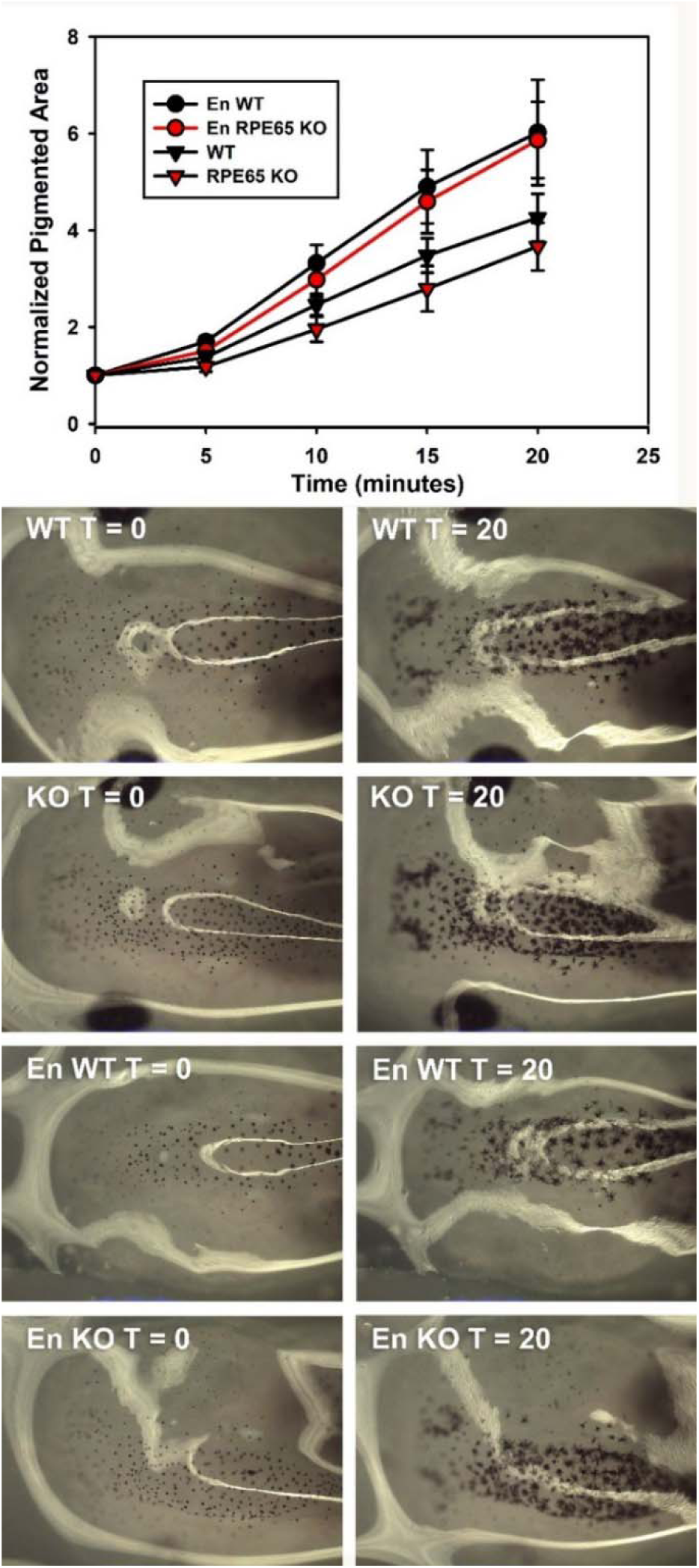
Pigment granule migration in tadpole melanocytes. WT, RPE65 KO, and enucleated WT and RPE65 KO animals were moved from dark to light condition, and photographed at several timepoints. Binary images were generated and the area of pigment was measured and plotted (top, representative images below). En = enucleated animal. See text for statistical analysis.

We found that melanophore dispersion continued over a period of at least 20 minutes under our assay conditions (Fig 6) (P (time) = 4.6E-14). As previously reported (Bertolesi et al., 2015; Hogben and Slome, 1936; Laurens, 1916), melanophore dispersion was influenced by the presence of eyes, progressing more rapidly in enucleated animals (P (enucleation) = 1.9E-5).

However, neither melanophore dispersion nor its alteration by enucleation were dependent on the presence of RPE65 (P (genotype) = 0.171, P (enucleation*genotype) = 0.65, P (time*genotype) = 0.98). Thus, neither the primary extraocular driver of melanophore dispersion in this assay, nor its moderation by the eye, requires RPE65.

## 4. Discussion

We previously generated *RPE65*-edited *X. laevis* for the purpose of investigating both the involvement of chromophore in retinal degeneration caused by *rhodopsin* mutations (Tam et al., 2024), and the involvement of chromophore in regulation of autophagy in rod photoreceptors (Wen et al., 2019). Here we have established a line of *X. laevis* carrying frameshift edits in all four *RPE65* alleles that completely prevent RPE65 expression, and characterized the resulting phenotype. Lack of expression is primarily due to knockout of *RPE65.L; RPE65.S* does not function significantly in the eyes of animals of the developmental stages examined. Notably, our results differ from those obtained with *rpe65* knockout mice in several interesting ways.

Our *RPE65* KO *X. laevis* retain all ciliated photoreceptors of the outer retina, including LWS and SWS cones, rods, and green rod photoreceptors. Green rods morphologically and functionally resemble rods but express a SWS cone pigment sensitive to blue wavelengths (Darden et al., 2003; Kojima et al., 2017). Histologically, the most notable difference between *RPE65* KO and WT *X. laevis* retinas was a decrease in the length of rods in older animals, similar to that seen in mice (Redmond et al., 1998). We did not find any evidence of mislocalized photopigments. This contrasts with *rpe65* knockout mice, in which SWS cones degenerate significantly by 2-3 weeks age (Znoiko et al., 2005), accompanied by inner segment mislocalization of cone opsin (Rohrer et al., 2005); the mechanism involves a biosynthetic defect in unliganded SWS cone opsin that is prevented by binding of chromophore (Zhang et al., 2011). In LCA patients, at least some cones are relatively long-lived, as their function can be measured by ERG in adult subjects (Jacobson et al., 2009), and gene therapy can restore some retinal function in adults (Bainbridge et al., 2015). Thus, *X. laevis* provide a useful system for investigating cone function in the absence of both RPE65 and retinal degeneration.

Like *rpe65* KO mice (Redmond et al., 1998), our *RPE65* KO *X. laevis* did not generate rhodopsin photopigment in sufficient quantities to assay by difference spectroscopy, and we were also unable to detect cone photopigments in difference spectra, although their abundance would be much lower than rhodopsin under all circumstances, and were not expected to be detectable. However, despite the lack of measurable photopigments, *RPE65* KO *X. laevis* eyes generate significant electrophysiological responses to light, with ∼2 log units loss in sensitivity relative to WT, indicating some photopigment(s) must be present; *rpe65* KO mice similarly have ERG responses, but lose >4 log units of sensitivity (Redmond et al., 1998) and human patients lose up to 5 log units of sensitivity by psychophysical measurements (Jacobson et al., 2007). In our older animals, both photopic and scotopic ERGs were easily measured at higher flash intensities. For a given flash intensity, a and b wave components were significantly delayed. In the case of scotopic ERGs, the responses of RPE65 KOs resembled WT responses to dimmer flashes. However, photopic ERG kinetics were more altered, suggesting that different cell types or physiological states may be involved. In contrast, mouse *rpe65* KO scotopic ERGs have altered (more rapid) kinetics, likely due to constitutive signaling via unliganded opsin inducing a physiological state similar to light adaptation (Fan et al., 2005). A possible explanation of this difference, as well as the lower reduction in sensitivity relative to mice, might be relatively lower constitutive signaling of rod opsin in *X. laevis*, but this requires further investigation.

Reduced photopic and scotopic ERG sensitivity occurred at all wavelengths tested, suggesting all *X. laevis* rods and cones have a significant requirement for RPE65. Several alternative pathways for photopigment generation in the absence of RPE65 have been suggested, and evidence for their existence has been obtained from different systems, including mice, frogs, and humans.

Our results show that RPE65-independent pathways confer small but significant ocular responses to a broad range of wavelengths of light in *X. laevis*. *X. laevis* photopigments primarily utilize 11-cis dehydroretinal (Bridges et al., 1977), resulting in a shift of ∼20nm in the absorption spectra relative to 11-cis retinal (Witkovsky et al., 1981). The complement of *X. laevis* retinal rod and cone photoreceptors includes a long-wavelength sensitive cone (575-620 nm depending on the report and chromophore), the dominant rod expressing the rhodopsin photopigment (500- 523 nm depending on chromophore), a SWS cone (425-445 nm depending on chromophore) and a rod-like photoreceptor (the green rod) which expresses a SWS cone pigment that absorbs at 434-445 nm depending on the chromophore (Bertolesi et al., 2022). Our ERG results suggest that long wavelength cones were relatively more affected than other cell types under scotopic conditions. As there is no medium-wavelength sensitive photopigment in *X. laevis*, our results also suggest that rods retain some function; this is similar to results with *rpe65* KO mice, in which residual rod activity is present, possibly driven by isorhodopsin, i.e. photopigment generated with 9-cis retinal (Fan et al., 2003). In future studies, we will compare these ERG data sets to similar data obtained with *X. laevis Rhodopsin* KOs currently under development.

We also looked for an effect on a visual behavior in *RPE65* KO *X. laevis*. We found that *RPE65* KO profoundly interfered with the ability of tadpoles to differentiate between light and dark backgrounds, indicating their vision is dramatically impaired. This response has been suggested to be mediated by SWS cones, and potentially a subset of SWS cones that express pinopsin in addition to the SWS1 pigment (Bertolesi et al., 2021). We also examined the response of melanophores to light; this response was not impaired, and neither was the specific component of this response that is regulated by the eye. The responses of melanophores to ambient light have been previously determined to be mediated by pineal photoreceptors maximally sensitive to green wavelengths, with additional regulation by melanopsin-expressing intrinsically photosensitive retinal ganglion cells that respond to substrate color (Bertolesi et al., 2021). Thus, our results indicate that neither pineal photoreceptors nor intrinsically photosensitive retinal ganglion cells require RPE65 for function in *X. laevis*.

Although several different pathways for RPE65-independent chromophore production have been proposed, there is controversy as to which of these pathways are relevant to visual function *in-vivo*. Our *X. laevis RPE65* KO animals will provide a useful system for assessing mechanisms of RPE65-independent responses via superimposed editing of additional genes proposed to function in these pathways, such as *DES1*, *MFAT*, and *RGR* opsin. As the *X. laevis* genome encodes representatives of all known vertebrate opsin gene classes, the utility of these animals may extend to the examination of opsins found in other retinal layers (such as melanopsin-expressing intrinsically photosensitive retinal ganglion cells) and/or non-retinal cell types.

## Acknowledgements

We would like to thank Dr. Masahiro Kono for the gift of the SWS2 (green rod) opsin antibody, as well as helpful discussions. This research was funded by the Canadian Institutes of Health Research (PJT-155937 and PJT-156072) and the National Science and Engineering Research Council (RGPIN-2020-05193).

## Author Contributions

Conceptualization: O. Moritz, B. Tam, C. Chiu; Experiments and data analysis: C. Chiu, B. Tam, O. Moritz, P. Burns; Writing and editing: C. Chiu: O. Moritz, B. Tam; Administration and securing funding: O. Moritz

## Supplementary Data

**Supplementary Table 1:**
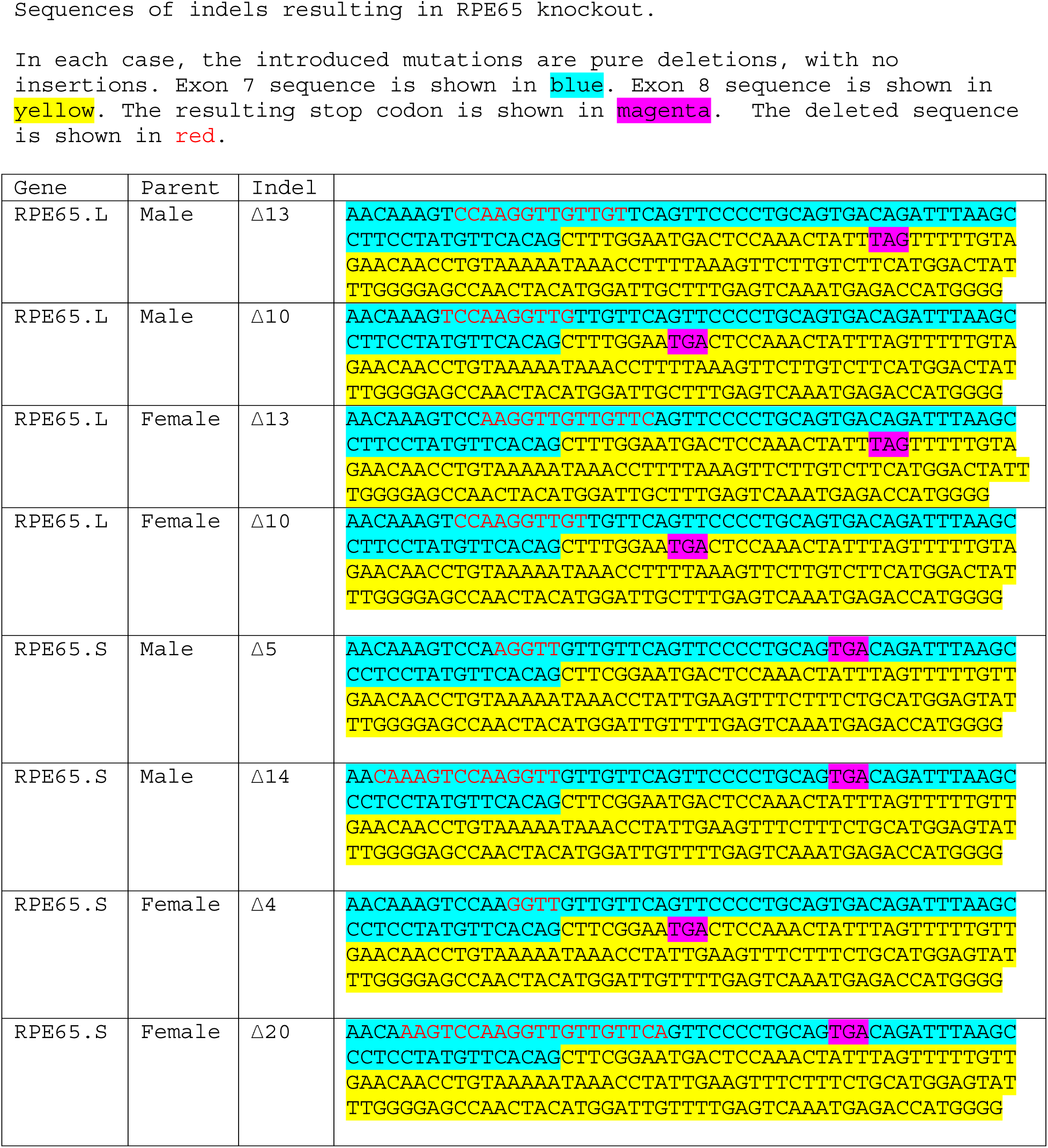
summary of indel mutations resulting in knockout:

## References

Acland, G.M., Aguirre, G.D., Bennett, J., Aleman, T.S., Cideciyan, A. V, Bennicelli, J., Dejneka, N.S., Pearce-Kelling, S.E., Maguire, A.M., Palczewski, K., Hauswirth, W.W., Jacobson, S.G., 2005. Long-term restoration of rod and cone vision by single dose rAAV-mediated gene transfer to the retina in a canine model of childhood blindness. Mol Ther 12, 1072– 1082. S1525-0016(05)01561-3 [pii]10.1016/j.ymthe.2005.08.008

Acland, G.M., Aguirre, G.D., Ray, J., Zhang, Q., Aleman, T.S., Cideciyan, A. V, Pearce-Kelling, S.E., Anand, V., Zeng, Y., Maguire, A.M., Jacobson, S.G., Hauswirth, W.W., Bennett, J., 2001. Gene therapy restores vision in a canine model of childhood blindness. Nat Genet 28, 92–95. 10.1038/8832788327 [pii]

Bainbridge, J.W.B., Mehat, M.S., Sundaram, V., Robbie, S.J., Barker, S.E., Ripamonti, C., Georgiadis, A., Mowat, F.M., Beattie, S.G., Gardner, P.J., Feathers, K.L., Luong, V.A., Yzer, S., Balaggan, K., Viswanathan, A., de Ravel, T.J.L., Casteels, I., Holder, G.E., Tyler, N., Fitzke, F.W., Weleber, R.G., Nardini, M., Moore, A.T., Thompson, D.A., Petersen-Jones, S.M., Michaelides, M., van den Born, L.I., Stockman, A., Smith, A.J., Rubin, G., Ali, R.R., 2015. Long-term effect of gene therapy on Leber’s congenital amaurosis. N. Engl. J. Med. 372, 1887–97. 10.1056/NEJMoa1414221

Bertolesi, G.E., Atkinson-Leadbeater, K., Mackey, E.M., Song, Y.N., Heyne, B., McFarlane, S., 2020. The regulation of skin pigmentation in response to environmental light by pineal Type II opsins and skin melanophore melatonin receptors. J. Photochem. Photobiol. B Biol. 212, 112024. 10.1016/J.JPHOTOBIOL.2020.112024

Bertolesi, G.E., Debnath, N., Atkinson-Leadbeater, K., Niedzwiecka, A., McFarlane, S., 2021. Distinct type II opsins in the eye decode light properties for background adaptation and behavioural background preference. Mol. Ecol. 30, 6659–6676. 10.1111/MEC.16203

Bertolesi, G.E., Debnath, N., Malik, H.R., Man, L.L.H., McFarlane, S., 2022. Type II Opsins in the Eye, the Pineal Complex and the Skin of Xenopus laevis: Using Changes in Skin Pigmentation as a Readout of Visual and Circadian Activity. Front. Neuroanat. 15, 118. 10.3389/FNANA.2021.784478/BIBTEX

Bertolesi, G.E., Hehr, C.L., Mcfarlane, S., 2015. Melanopsin photoreception in the eye regulates light-induced skin colour changes through the production of α-MSH in the pituitary gland. Pigment Cell Melanoma Res. 28, 559–571. 10.1111/PCMR.12387

Bridges, C.D.B., Hollyfield, J.G., Witkovsky, P., Gallin, E., 1977. The visual pigment and vitamin A of Xenopus laevis embryos, larvae and adults. Exp. Eye Res. 24, 7–13. 10.1016/0014-4835(77)90279-2

Bruch Goldstein, E., Wolf, B.M., 1973. Regeneration of the green-rod pigment in the isolated frog retina. Vision Res. 13, 527–534. 10.1016/0042-6989(73)90022-9

Cai, X., Conley, S.M., Naash, M.I., 2009. RPE65: Role in the visual cycle, human retinal disease, and gene therapy. Ophthalmic Genet. 10.1080/13816810802626399

Chen, P., Hao, W., Rife, L., Wang, X.P., Shen, D., Chen, J., Ogden, T., Van Boemel, G.B., Wu, L., Yang, M., Fong, H.K.W., 2001. A photic visual cycle of rhodopsin regeneration is dependent on Rgr. Nat. Genet. 28, 256–260. 10.1038/90089

Cideciyan, A. V, Aleman, T.S., Boye, S.L., Schwartz, S.B., Kaushal, S., Roman, A.J., Pang, J.J., Sumaroka, A., Windsor, E.A., Wilson, J.M., Flotte, T.R., Fishman, G.A., Heon, E., Stone, E.M., Byrne, B.J., Jacobson, S.G., Hauswirth, W.W., 2008. Human gene therapy for RPE65 isomerase deficiency activates the retinoid cycle of vision but with slow rod kinetics. Proc Natl Acad Sci U S A 105, 15112–15117. 0807027105<otherinfo> [pii]</otherinfo>10.1073/pnas.0807027105

Darden, A.G., Wu, B.X., Znoiko, S.L., Hazard, E.S., Kono, M., Crouch, R., Ma, J.-X., 2003. Darden, Mol Vis 2003; 9:191–199. Mol. Vis. 9, 191–199.

Fan, J., Rohrer, B., Moiseyev, G., Ma, J.X., Crouch, R.K., 2003. Isorhodopsin rather than rhodopsin mediates rod function in RPE65 knock-out mice. Proc. Natl. Acad. Sci. U. S. A. 100, 13662–13667. 10.1073/PNAS.2234461100/ASSET/051A89A2-4C1C-4F8F-9C81-3290DCBD43F5/ASSETS/GRAPHIC/PQ2234461005.JPEG

Fan, J., Woodruff, M.L., Cilluffo, M.C., Crouch, R.K., Fain, G.L., 2005. Opsin activation of transduction in the rods of dark-reared Rpe65 knockout mice. J. Physiol. 568, 83–95. 10.1113/JPHYSIOL.2005.091942

Feehan, J.M., Chiu, C.N., Stanar, P., Tam, B.M., Ahmed, S.N., Moritz, O.L., 2017. Modeling Dominant and Recessive Forms of Retinitis Pigmentosa by Editing Three Rhodopsin-Encoding Genes in Xenopus Laevis Using Crispr/Cas9. Sci. Rep. 7. 10.1038/s41598-017-07153-4

Feehan, J.M., Stanar, P., Tam, B.M., Chiu, C., Moritz, O.L., 2019. Generation and Analysis of Xenopus laevis Models of Retinal Degeneration Using CRISPR/Cas9, in: Methods in Molecular Biology (Clifton, N.J.). pp. 193–207. 10.1007/978-1-4939-8669-9_14

Ghaseminejad, F., Tam, B.M., Chiu, C.N., Feehan, J.M., Moritz, O.L., 2022. Gene editing treatment strategies for retinitis pigmentosa assessed in Xenopus laevis carrying a mutant Rhodopsin allele. J. Transl. Genet. Genomics 6, 111–125. 10.20517/JTGG.2021.49

Goldstein, E.B., 1970. Cone pigment regeneration in the isolated frog retina. Vision Res. 10, 1065–1068. 10.1016/0042-6989(70)90082-9

Hogben, L., Slome, D., 1936. The pigmentary effector system VIII—The dual receptive mechanism of the amphibian background response. Proc. R. Soc. London. Ser. B - Biol. Sci. 120, 158–173. 10.1098/RSPB.1936.0029

Jacobson, S.G., Aleman, T.S., Cideciyan, A. V., Roman, A.J., Sumaroka, A., Windsor, E.A.M.M., Schwartz, S.B., Heon, E., Stone, E.M., 2009. Defining the residual vision in leber congenital amaurosis caused by RPE65 mutations. Invest Ophthalmol Vis Sci 50, 2368–2375. iovs.08-2696 [pii]10.1167/iovs.08-2696

Jacobson, S.G., Aleman, T.S., Cideciyan, A. V, Heon, E., Golczak, M., Beltran, W.A., Sumaroka, A., Schwartz, S.B., Roman, A.J., Windsor, E.A., Wilson, J.M., Aguirre, G.D., Stone, E.M., Palczewski, K., 2007. Human cone photoreceptor dependence on RPE65 isomerase. Proc Natl Acad Sci U S A 104, 15123–15128. 0706367104 [pii]10.1073/pnas.0706367104

Jin, M., Li, S., Moghrabi, W.N., Sun, H., Travis, G.H., 2005. Rpe65 is the retinoid isomerase in bovine retinal pigment epithelium. Cell 122, 449–59. 10.1016/j.cell.2005.06.042

Kaylor, J.J., Cook, J.D., Makshanoff, J., Bischoff, N., Yong, J., Travis, G.H., 2014. Identification of the 11-cis-specific retinyl-ester synthase in retinal Müller cells as multifunctional O-Acyltransferase (MFAT). Proc. Natl. Acad. Sci. U. S. A. 111, 7302–7307. 10.1073/PNAS.1319142111/-/DCSUPPLEMENTAL/PNAS.201319142SI.PDF

Kaylor, J.J., Xu, T., Ingram, N.T., Tsan, A., Hakobyan, H., Fain, G.L., Travis, G.H., 2017. Blue light regenerates functional visual pigments in mammals through a retinyl-phospholipid intermediate. Nat. Commun. 8, 16. 10.1038/s41467-017-00018-4

Kaylor, J.J., Yuan, Q., Cook, J., Sarfare, S., Makshanoff, J., Miu, A., Kim, A., Kim, P., Habib, S., Roybal, C.N., Xu, T., Nusinowitz, S., Travis, G.H., 2013. Identification of DES1 as a vitamin A isomerase in Müller glial cells of the retina. Nat. Chem. Biol. 9, 30–36. 10.1038/nchembio.1114

Kiser, P.D., Kolesnikov, A. V., Kiser, J.Z., Dong, Z., Chaurasia, B., Wang, L., Summers, S.A., Hoang, T., Blackshaw, S., Peachey, N.S., Kefalov, V.J., Palczewski, K., 2019. Conditional deletion of Des1 in the mouse retina does not impair the visual cycle in cones. FASEB J. 33, 5782–5792. 10.1096/FJ.201802493R

Kojima, K., Matsutani, Y., Yamashita, T., Yanagawa, M., Imamoto, Y., Yamano, Y., Wada, A., Hisatomi, O., Nishikawa, K., Sakurai, K., Shichida, Y., 2017. Adaptation of cone pigments found in green rods for scotopic vision through a single amino acid mutation. Proc. Natl. Acad. Sci. U. S. A. 114, 5437–5442. 10.1073/PNAS.1620010114/SUPPL_FILE/PNAS.201620010SI.PDF

Laurens, H., 1916. The reactions of the melanophores of Amblystoma larvae—the supposed influence of the pineal organ. J. Exp. Zool. 20, 237–261. 10.1002/JEZ.1400200208

Ma, J. xing, Xu, L., Othersen, D.K., Redmond, T.M., Crouch, R.K., 1998. Cloning and localization of RPE65 mRNA in salamander cone photoreceptor cells1. Biochim. Biophys. Acta 1443, 255–261. 10.1016/S0167-4781(98)00221-8

Marlhens, F., Bareil, C., Griffoin, J.M., Zrenner, E., Amalric, P., Eliaou, C., Liu, S.Y., Harris, E., Redmond, T.M., Arnaud, B., Claustres, M., Hamel, C.P., 1997. Mutations in RPE65 cause Leber’s congenital amaurosis. Nat. Genet. 17, 139–41. 10.1038/ng1097-139

Moiseyev, G., Chen, Y., Takahashi, Y., Wu, B.X., Ma, J.-X., 2005. RPE65 is the isomerohydrolase in the retinoid visual cycle. Proc. Natl. Acad. Sci. U. S. A. 102, 12413–8. 10.1073/pnas.0503460102

Molday, R.S., Moritz, O.L., 2015. Photoreceptors at a glance. J. Cell Sci. 128. 10.1242/jcs.175687

Montell, C., 2012. Drosophila visual transduction. Trends Neurosci. 35, 356. 10.1016/J.TINS.2012.03.004

Moriya, T., Kito, K., Miyashita, Y., Asami, K., 1996. Preference for Background Color of the Xenopus Zaevis Tadpole. J. Exp. Zool. 276–335. 10.1002/(SICI)1097-010X(19961201)276:5

Morshedian, A., Kaylor, J.J., Ng, S.Y., Tsan, A., Frederiksen, R., Xu, T., Yuan, L., Sampath, A.P., Radu, R.A., Fain, G.L., Travis, G.H., 2019. Light-Driven Regeneration of Cone Visual Pigments through a Mechanism Involving RGR Opsin in Müller Glial Cells. Neuron 102, 1172–1183.e5. 10.1016/j.neuron.2019.04.004

Redmond, T.M., Poliakov, E., Yu, S., Tsai, J.Y., Lu, Z., Gentleman, S., 2005. Mutation of key residues of RPE65 abolishes its enzymatic role as isomerohydrolase in the visual cycle. Proc. Natl. Acad. Sci. U. S. A. 102, 13658–13663. 10.1073/PNAS.0504167102/ASSET/52FA8098-3B85-47DD-B80F-A1B120CD4E71/ASSETS/GRAPHIC/ZPQ0380595490006.JPEG

Redmond, T.M., Yu, S., Lee, E., Bok, D., Hamasaki, D., Chen, N., Goletz, P., Ma, J.X., Crouch, R.K., Pfeifer, K., 1998. Rpe65 is necessary for production of 11-cis-vitamin A in the retinal visual cycle. Nat. Genet. 20, 344–51. 10.1038/3813

Rohrer, B., Lohr, H.R., Humphries, P., Redmond, T.M., Seeliger, M.W., Crouch, R.K., 2005. Cone opsin mislocalization in Rpe65-/- mice: a defect that can be corrected by 11-cis retinal. Invest. Ophthalmol. Vis. Sci. 46, 3876–3882. 10.1167/IOVS.05-0533

Saari, J.C., 2000. Biochemistry of visual pigment regeneration: the Friedenwald lecture. Invest. Ophthalmol. Vis. Sci. 41, 337–48.

Session, A.M., Uno, Y., Kwon, T., Chapman, J.A., Toyoda, A., Takahashi, S., Fukui, A., Hikosaka, A., Suzuki, A., Kondo, M., van Heeringen, S.J., Quigley, I., Heinz, S., Ogino, H., Ochi, H., Hellsten, U., Lyons, J.B., Simakov, O., Putnam, N., Stites, J., Kuroki, Y., Tanaka, T., Michiue, T., Watanabe, M., Bogdanovic, O., Lister, R., Georgiou, G., Paranjpe, S.S., van Kruijsbergen, I., Shu, S., Carlson, J., Kinoshita, T., Ohta, Y., Mawaribuchi, S., Jenkins, J., Grimwood, J., Schmutz, J., Mitros, T., Mozaffari, S. V., Suzuki, Y., Haramoto, Y., Yamamoto, T.S., Takagi, C., Heald, R., Miller, K., Haudenschild, C., Kitzman, J., Nakayama, T., Izutsu, Y., Robert, J., Fortriede, J., Burns, K., Lotay, V., Karimi, K., Yasuoka, Y., Dichmann, D.S., Flajnik, M.F., Houston, D.W., Shendure, J., DuPasquier, L., Vize, P.D., Zorn, A.M., Ito, M., Marcotte, E.M., Wallingford, J.B., Ito, Y., Asashima, M., Ueno, N., Matsuda, Y., Veenstra, G.J.C., Fujiyama, A., Harland, R.M., Taira, M., Rokhsar, D.S., 2016. Genome evolution in the allotetraploid frog Xenopus laevis. Nature 538, 336–343. 10.1038/nature19840

Stavenga, D.G., 1995. Insect retinal pigments: Spectral characteristics and physiological functions. Prog. Retin. Eye Res. 15, 231–259. 10.1016/1350-9462(95)00011-9

Tam, B.M., Paloma, B., Chiu, C.N., Moritz, O.L., 2024. Synchronized photoactivation of T4K rhodopsin causes a chromophore-dependent retinal degeneration that is moderated by interaction with phototransduction cascade components. J. Neurosci. e0453242024. 10.1523/JNEUROSCI.0453-24.2024

Tworak, A., Kolesnikov, A. V., Hong, J.D., Choi, E.H., Luu, J.C., Palczewska, G., Dong, Z., Lewandowski, D., Brooks, M.J., Campello, L., Swaroop, A., Kiser, P.D., Kefalov, V.J., Palczewski, K., 2023. Rapid RGR-dependent visual pigment recycling is mediated by the RPE and specialized Müller glia. Cell Rep. 42, 112982. 10.1016/J.CELREP.2023.112982

Veske, A., Nilsson, S.E.G., Narfström, K., Gal, A., 1999. Retinal dystrophy of Swedish briard/briard-beagle dogs is due to a 4-bp deletion in RPE65. Genomics 57, 57–61. 10.1006/GENO.1999.5754

Viczian, A.S., Zuber, M.E., 2014. A simple behavioral assay for testing visual function in Xenopus laevis. J. Vis. Exp. 10.3791/51726

Wang, J.S., Kefalov, V.J., 2011. The cone-specific visual cycle. Prog. Retin. Eye Res. 30, 115–128. 10.1016/J.PRETEYERES.2010.11.001

Ward, R., Kaylor, J.J., Cobice, D.F., Pepe, D.A., McGarrigle, E.M., Brockerhoff, S.E., Hurley, J.B., Travis, X.G.H., Kennedy, X.B.N., 2020. Non-photopic and photopic visual cycles differentially regulate immediate, early, and late phases of cone photoreceptor-mediated vision. J. Biol. Chem. 295, 6482–6497. 10.1074/JBC.RA119.011374

Wen, R.H., Stanar, P., Tam, B., Moritz, O.L., 2019. Autophagy in Xenopus laevis rod photoreceptors is independently regulated by phototransduction and misfolded RHOP23H. Autophagy 1–20. 10.1080/15548627.2019.1596487

Witkovsky, P., Levine, J.S., Engbretson, G.A., Hassin, G., MacNichol Jr., E.F., 1981. A microspectrophotometric study of normal and artificial visual pigments in the photoreceptors of Xenopus laevis. Vis. Res 21, 867–873.

Zhang, T., Zhang, N., Baehr, W., Fu, Y., 2011. Cone opsin determines the time course of cone photoreceptor degeneration in Leber congenital amaurosis. Proc. Natl. Acad. Sci. U. S. A. 108, 8879–8884. 10.1073/PNAS.1017127108/-/DCSUPPLEMENTAL/SAPP.PDF

Znoiko, S.L., Crouch, R.K., Moiseyev, G., Ma, J.X., 2002. Identification of the RPE65 protein in mammalian cone photoreceptors. Investig. Ophthalmol. Vis. Sci. 43, 1604–1609.

Znoiko, S.L., Rohrer, B., Lu, K., Lohr, H.R., Crouch, R.K., Ma, J.X., 2005. Downregulation of cone-specific gene expression and degeneration of cone photoreceptors in the Rpe65-/- mouse at early ages. Invest. Ophthalmol. Vis. Sci. 46, 1473–1479. 10.1167/IOVS.04-0653

